# EBV Reprograms B Cells in an Autoimmune-Like Fashion in Patients with COVID-19

**DOI:** 10.64898/2026.08.17.745245

**Authors:** Daniel G. Chen, Dan Yuan, Yapeng Su, Andrew Magis, Helen Chu, Jason D. Goldman, James R. Heath

**Affiliations:** UCLA-Caltech Medical Scientist Training Program, University of California, Los Angeles, Los Angeles, CA; Institute of Systems Biology, Seattle, WA; Fred Hutchinson Cancer Center, Seattle, WA; Division of Global Health, University of Washington, Seattle, WA; Division of Allergy and Infectious Diseases, Department of Medicine, University of Washington, Seattle, WA; Swedish Center for Research and Innovation, Providence Swedish Medical Center, Seattle, WA; Department of Bioengineering, University of Washington, Seattle, WA

## Abstract

Epstein-Barr virus (EBV) reprograms B cells in autoimmune disease. Reprogrammed EBV^+^ B cells activate nearby B and CD4^+^ T cells, via upregulated antigen presentation and costimulatory machinery, to drive autoimmune pathology. EBV reactivation is a known correlate of long COVID, which is a heterogeneous condition that can bear similarities to autoimmune disease. However, the mechanisms underpinning this association remain unresolved. We report on EBV metabolically reprogrammed B cells in patients with COVID-19. We find EBV^+^ B cells provide stimulatory signals to bystander B and CD4^+^ T cells. SARS-CoV-2 infected participants exhibiting elevated fractions of EBV^+^ B cells present, at convalescence, with dysregulated lipid profiles, increased autoantibody titers, and post-acute symptomology likely reflective of this metabolic reprogramming and cell-cell interactions. Enrichment of our EBV^+^ B cell signatures seen in patients with COVID-19 is similar in patients with lupus and multiple sclerosis suggesting a potentially shared pathway of EBV-driven dysfunction across diseases.

## Introduction

Epstein-Barr virus (EBV) is a human gamma herpesvirus that establishes lifelong latency with intermittent reactivation to lytic replication. EBV causes infectious mononucleosis^1^ in a minority with acute infection, and is a well-known oncogenic virus, associated with multiple cancers (e.g. Burkitt lymphoma^2^, gastric cancer^3^ and others^4^), and is reported to have infected ≥90% of the global population. In the past decade, large-scale studies have demonstrated a consistent positive association between prior EBV infection and increased autoimmune disease risk, particularly for multiple sclerosis (MS)^5–7^. These efforts have proposed molecular mimicry (similarity between human and EBV antigens) and EBV transcription factor (e.g. EBNA2) activity as two potential mechanisms driving EBV-associated autoimmune disease pathology^8–10^. Recent works have compounded on these findings by demonstrating a fundamental reprogramming of EBV-infected B cells towards a metabolically-distinct cell state^11–14^ (e.g. increased glycolysis) that markedly upregulates antigen presentation and co-stimulation machinery^15–18^. Reprogrammed EBV-infected B cells can then leverage this machinery to activate autoreactive B and CD4^+^ T cells and thus facilitate autoimmune disease pathology, such as autoantibody production.

Long COVID is a debilitating post-acute sequelae of SARS-CoV-2 infection that continues to affect hundreds of millions of people worldwide. An estimated 7% of adults in the United States alone have had long COVID (as reported in 2024), with global numbers estimated as even higher ^19,20^. Patients with long COVID can be afflicted with a range of symptoms, including brain fog, post-exertional malaise, chronic fatigue, respiratory, digestive, and/or cardiovascular issues that can severely limit their quality of life^19,21,22^. To address this patient hardship, many studies have sought to identify long COVID risk factors shared across patient populations, which may thus represent potential therapeutic targets. These studies have consistently revealed a marked positive association between long COVID risk and EBV reactivation (or detection via viremia)^23–28;^ these findings have been increasingly validated in larger patient populations such as the IMPACC cohort^29^. While these works establish a clear associative link between EBV and long COVID, they do not identify underpinning molecular mechanisms. The similarity between certain immunopathology in long COVID (e.g. dysregulated T cells and B cells and autoantibodies^30^) and recently identified EBV driven mechanisms of disease in systemic lupus erythematous (SLE) and MS raise the possibility of EBV infection as not only a correlate of long COVID but an active effector that drives the condition’s pathology in at least a fraction of patients.

## Results

### Identification of EBV^+^ B cells in patients with COVID-19

We leveraged a reported computational framework in autoimmune disease^16,17^ to identify EBV-infected B cells in a primary cohort of 209 patients with COVID-19 (INCOV) and a validation cohort of 100 patients with COVID-19 (HAARVI)^25^ (**Fig. 1A, Methods**); the INCOV cohort was used for the majority of analyses unless otherwise specified.

**Fig. 1:**
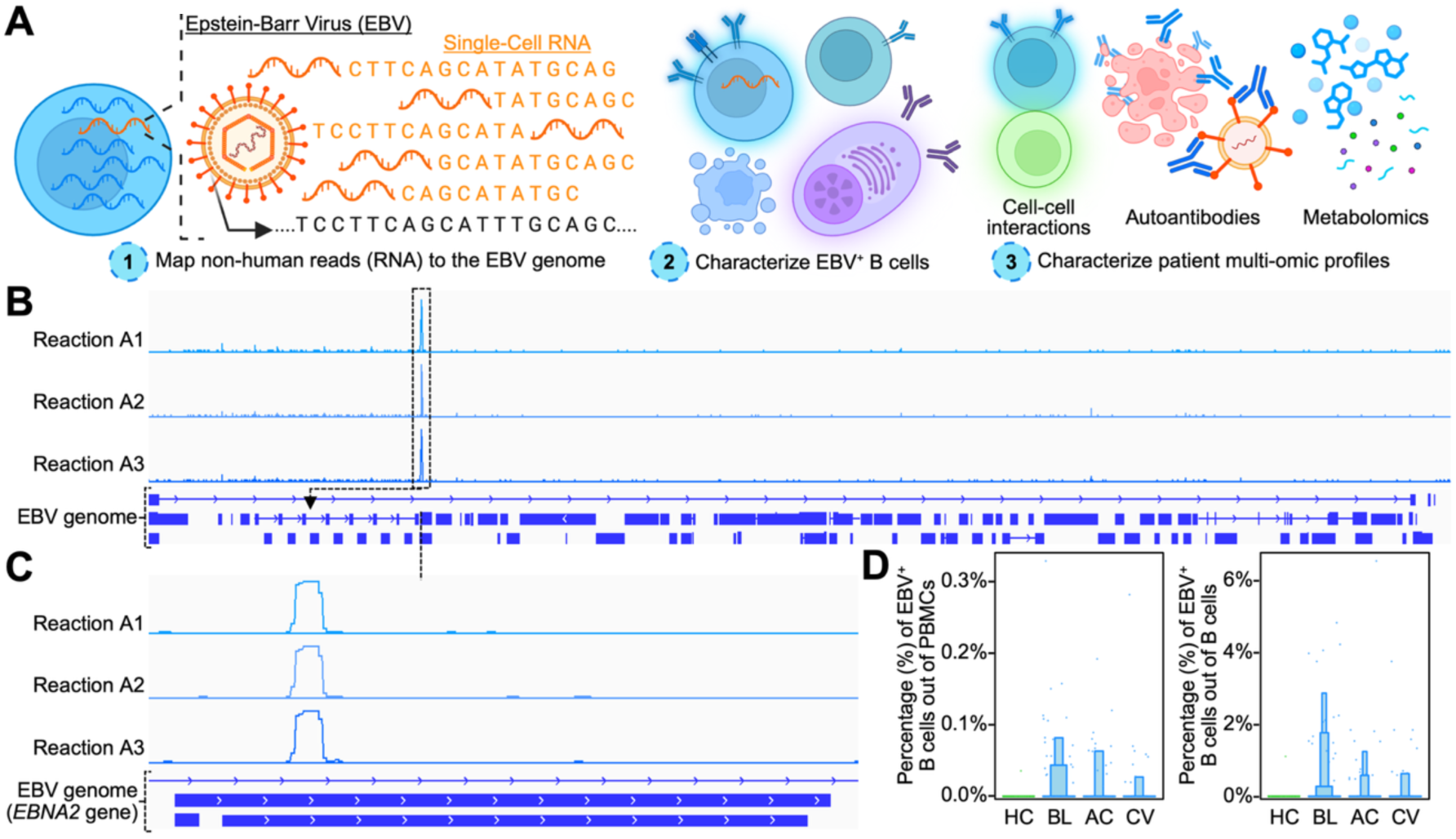
Computational identification of EBV^+^ cells in PBMCs from patients with COVID-19. A) Diagram of the EBV read identification process (1), whereby reads that did not map to the human genome were re-mapped against the EBV genome. EBV^+^ B cells were then characterized via scRNAseq and metabolic flux analysis (2). Patients with high or low proportions of EBV^+^ B cells were then subjected to comparisons against plasma metabolites or autoantibody levels. B) Track plots of reads mapped to the EBV genome (NC_007605.1) across three representative 10x reactions (A1, A2, A3). The entire EBV genome is presented below the three track plots with exons represented by solid blue blocks. C) Track plots of reads mapped to the *EBNA2* gene from the EBV genome (a transcriptional activator expressed during latency III) across three representative 10x reactions. Exons are represented by solid blue blocks. D) Boxen plots of the percentage of a given sample (blood draw) with detectable EBV reads out of all PBMCs (left) or B cells (right). Samples are stratified by disease state: healthy donors (HD), baseline (BL, diagnosis of COVID-19), acute disease (AC, one week after diagnosis), and convalescence (CV, 2-3 months after acute disease). Boxen plots represent interquartile range (25 to 75 percentiles) with successive boxes representing progressively halved quantiles (e.g. 12.5/87.5, 6.25/93.75 percentiles). Dots represent individual samples.

Detected EBV reads from single-cell RNA-seq (scRNAseq) of peripheral mononuclear blood cells (PBMCs), largely mapped to the *EBNA2* gene (**Figs. 1B-C**). *EBNA2* is a well-known transcriptional activator expressed during latency III of EBV infection^31^ and reported to bind at transcription-start-sites of genes involved in B cell survival, activation, and antigen presentation^16,32,33^. EBV^+^ B cells were initially identified using a strict cutoff for reads that uniquely mapped to the EBV genome (**Fig. 1D**); this revealed an average of 0.17±0.06% EBV^+^ B cells. As viruses bear highly repetitive genomes^34^, viral reads often map multiple times to their genome and are thus lost when using strict cutoffs that require unique mapping. To overcome this, we also assigned B cells with viral reads that mapped multiple times to the EBV genome as EBV^+^. We confirmed the validity of this approach by finding all EBV^+^ B cells to enrich for an EBV-specific gene signature regardless of the number of times their EBV reads mapped to the EBV genome (**Fig. S1**). This resulted in an average of 2.76±0.27% EBV^+^ B cells across patients and healthy donors (**Table S1**). These EBV^+^ assignments were subsequently utilized for downstream analyses. Notably, few EBV^+^ B cells could be detected in healthy donors in comparison with patients with COVID-19. This finding is consistent with prior reports of EBV reactivation after SARS-CoV-2 infection^16,23,24,26,27^ and focused our subsequent analyses on the EBV^+^ B cells derived from patients with COVID-19.

### EBV transcriptionally reprograms B cells in patients with COVID-19

Inspired by recent reports of marked B cell reprogramming by EBV in autoimmune disease^16–18^, we hypothesized that EBV^+^ B cells may undergo a similar reprogramming in patients with COVID-19. To investigate this, we first classified single B cells into canonical B cell states by their marker gene expression (e.g. *FCER2* in naïve B cells^35^) (**Figs. 2A and S2A**). We then interrogated for differences in the distribution of B cell states in EBV^+^ versus EBV^-^ B cell compartments. This revealed EBV^+^ B cells to be significantly biased towards memory B cell and plasmablast states (**Fig. 2B**) and is reminiscent of previous findings of EBV-transformed B cells as highly proliferative and plasmablast-like^36^. EBV^+^ naïve B cells suggests recent infection from viral particles released by EBV-infected B cells that have undergone completion of a reactivated lytic cycle^37^.

**Fig. 2:**
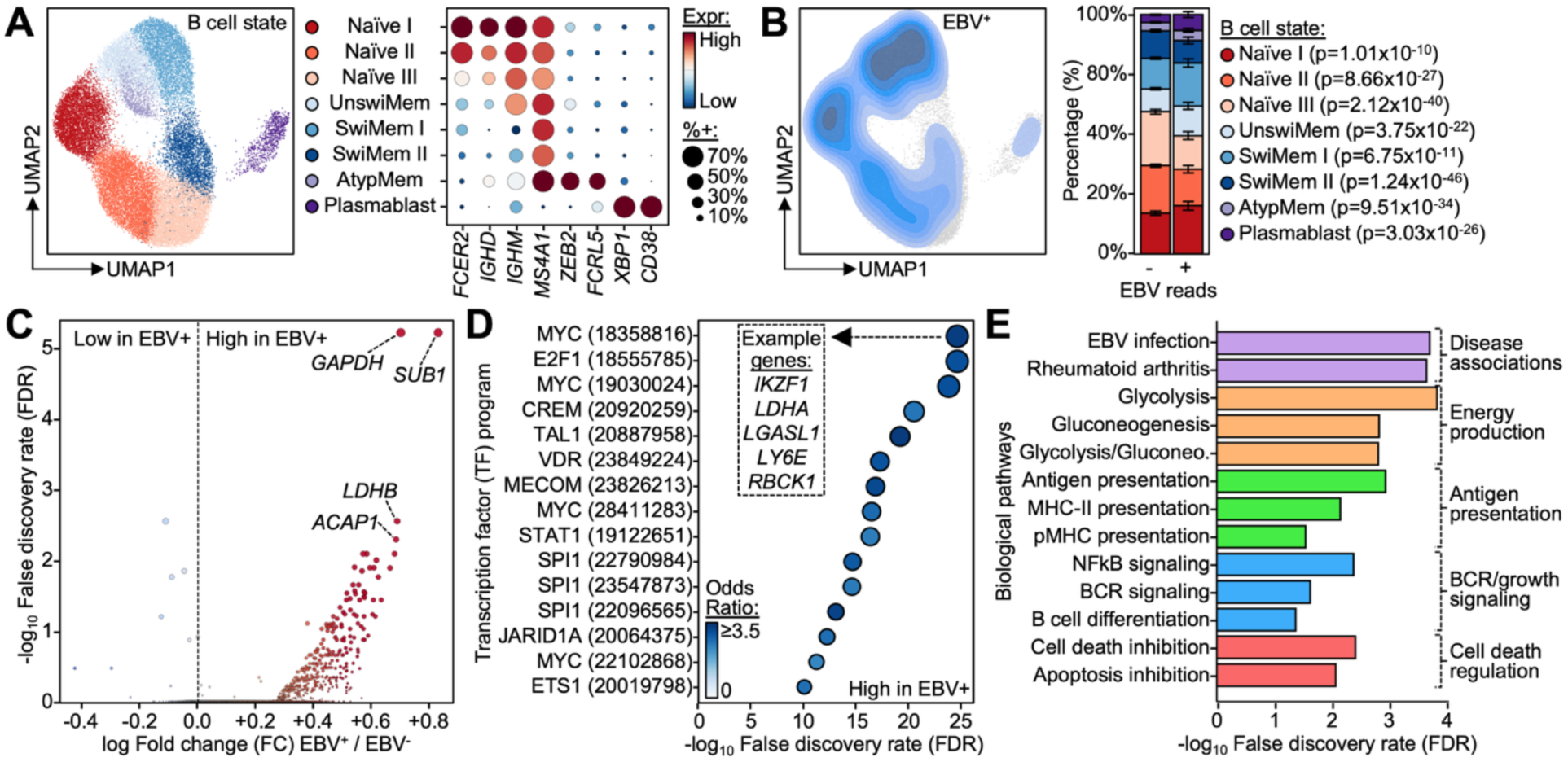
EBV reprograms B cells towards distinct transcriptional cell states in COVID-19. A) Left: Uniform manifold approximation projection (UMAP) of single B cell transcriptomes. Cells are colored by their cell state (see legend on the right); unswitched memory is abbreviated as UnswiMem, switched memory as SwiMem, and atypical memory as AtypMem. Right: Dot plot of the expression of B cell marker genes (columns) by cell state (rows). Dot size represents the percent of a given cell state that expresses said marker gene and color represents expression magnitude (see legend on the right). B) Left: Contour plot of the distribution of EBV^+^ B cells on the UMAP from panel (A). Right: Stacked bar plots of the distribution of B cell states amongst EBV^-^ and EBV^+^ B cells (left and right bars, respectively). Bars are colored by cell state (see legend on the right). Bar height represents arithmetic mean, and error bars represent standard error. P-values were calculated using the Mann-Whitney U test (right). C) Volcano plot of the differentially expressed genes in EBV^+^ B cells compared to EBV^-^ B cells. X-axis and color represent log-transformed fold change in gene expression (genes upregulated in EBV^+^ B cells are on the right). Y-axis represents the significance of the difference, -log_10_ false discovery rate (FDR). D) Dot plot of the enrichment of transcription factor programs (from ChEA, on the Y-axis) as quantified by -log_10_(FDR) (represented on the X-axis and by dot size). Color represents odds ratio (see legend on the lower left). Numbers on the Y-axis denote dataset (PubMed) ID. E) Bar plot of the pathways enriched in the differentially expressed genes from panel (C) in EBV^+^ B cells (on the Y-axis) as quantified by -log_10_(FDR) (on the Y-axis). Color represents shared pathway categories (e.g. antigen presentation).

To identify the exact transcriptional modules reprogrammed by EBV, we performed differential gene expression analysis between EBV^+^ and EBV^-^ B cells (**Fig. 2C**, **Table S2**). Interestingly, we observed EBV^+^ B cells to present with more significantly upregulated (N=30, FDR<0.05) rather than downregulated genes (N=3, FDR<0.05). This is consistent with the expression of the EBV-derived transcriptional activator *EBNA2* in EBV^+^ B cells. Transcription factor (TF) program enrichment analysis revealed EBV^+^ B cells to activate well-known EBV-controlled TFs (**Figs. 2D and S2B, Table S3**), such as MYC and SPI1 which drive B cell proliferation and interact with EBV proteins (e.g. EBNA2)^38–40^. Pathway enrichment analysis revealed a distinct reprogramming of EBV^+^ B cells towards increased antigen presentation, energy production via glycolysis, and B cell activation and survival signaling (**Fig. 2E, Table S4**). Consistently, EBV has been reported to upregulate glycolytic^11,41^ and anti-apoptotic functions^42,43^ and drives increased antigen presentation in autoimmune disease^16,18^. Increased B cell activation in EBV^+^ B cells was confirmed by kinase enrichment analysis revealing increased BTK, SYK, and ZAP70 activity^44–46^ (**Table S5**). In conjunction with the enrichment of EBV infection associated genes in EBV^+^ B cells, these data suggest that patients with COVID-19 experience *bona fide* EBV reprogramming of their B cells towards a distinct transcriptional cell state. We confirmed these findings of EBV reprogrammed B cells in our independent validation cohort of 100 patients with COVID-19 (HAARVI) (**Figs. S2C-D**).

### EBV reprogrammed B cells in COVID-19 resemble those in autoimmune disease

The enrichment of antigen presentation and B cell activation processes in EBV reprogrammed B cells in the context of COVID-19 is reminiscent of the EBV reprogramming observed in certain autoimmune diseases^16,18^. This is supported by the significant enrichment of autoimmune disease related genes (e.g. SLE) in EBV^+^ B cells (**Table S10**). To formally investigate this similarity, we queried for the expression of genes (e.g. *CD27*, *CD70*, *IL2RG*, *IGHA1, PTPN22*) and functions (e.g. proteasome, antigen presentation, interferon and B cell activation signaling) reported to be specifically upregulated in EBV reprogrammed B cells in autoimmune disease^11,16–18^. These markers were, by and large, all significantly upregulated in EBV^+^ B cells in patients with COVID-19 (**Figs. 3A and S3A**). We confirmed these observations by finding EBV^+^ B cells in patients with COVID-19 to enrich for reported transcriptomic signatures of EBV^+^ B cells in SLE^16^ (**Fig. 3B**). To deepen our understanding of these cells, we repeated the above analyses in our recently reported multiome (transcriptome and epigenome from the same single cell) dataset of B cells from patients with COVID-19 (**Fig. S3B**)^30^. In confirmation of our previous findings, identified EBV^+^ B cells once again enriched for cell populations (e.g. IgA^+^ plasmablasts, **Fig. S3C**) and gene signatures (**Fig. S3D**) reported to be associated with EBV^+^ B cells in autoimmune contexts^16–18^. Epigenetic analysis of differentially open chromatin in EBV^+^ B cells revealed significant co-enrichment for RBPJ and EBF1 transcription factor (TF) binding sites (**Fig. S3E**); these shared binding sites were enriched for EBV associated genes, e.g. *PTPN6*^47^ (**Fig. S3F**). RBPJ and EBF1 are known to be co-opted by the EBV protein EBNA2 to form a transcriptional complex that drives the pathogenic transformation of EBV^+^ B cells^48^. Further, these TFs have been reported to have increased activity in EBV^+^ B cells from patients with autoimmune disease^16^. Together, these data suggest EBV^+^ B cells in patients with COVID-19 to have undergone canonical EBV reprogramming in terms of their transcriptomes (individual genes and pathways) and epigenomes, similar to EBV^+^ B cells in certain autoimmune diseases.

**Fig. 3:**
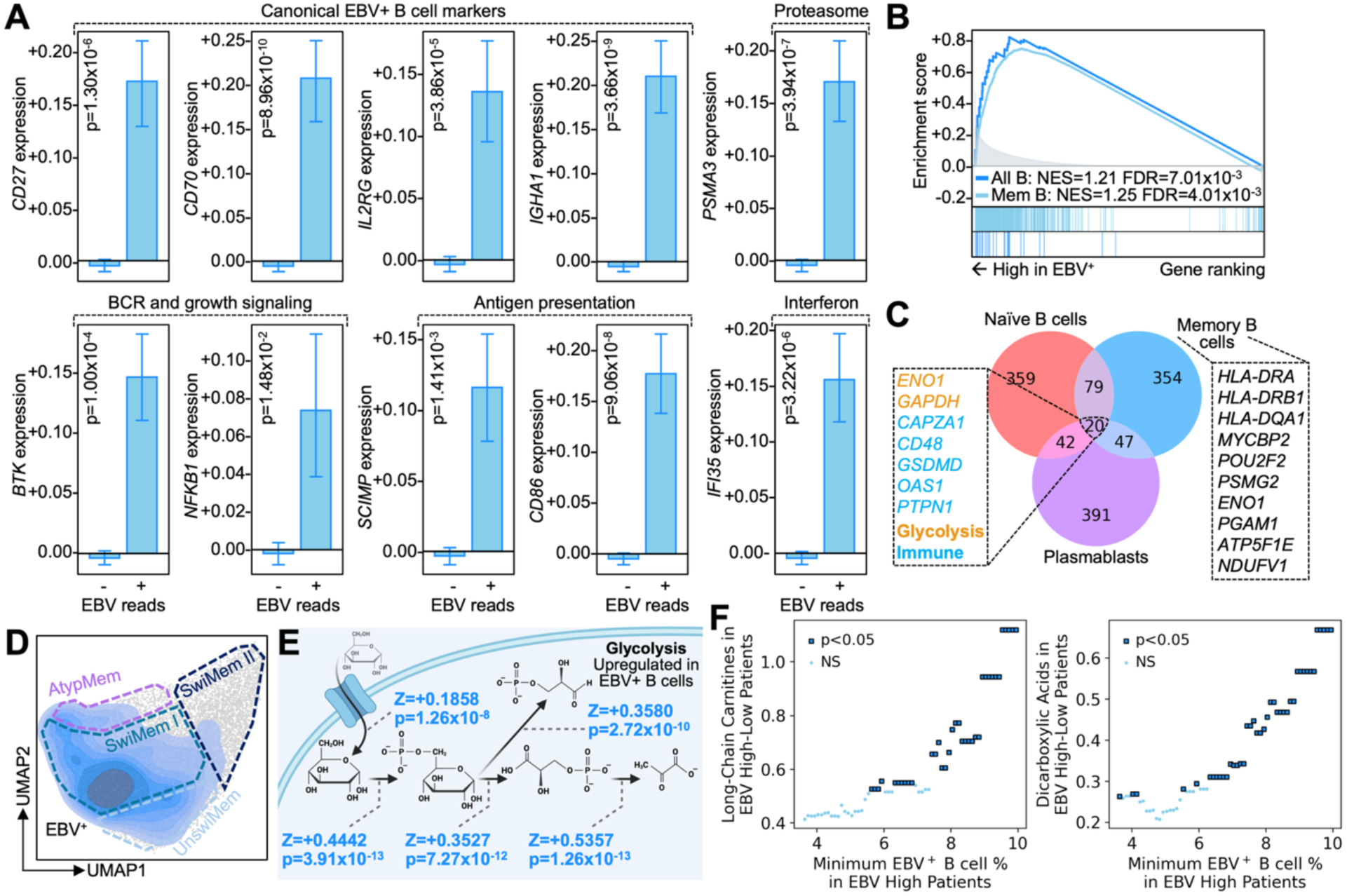
EBV reprogrammed B cells in COVID-19 are metabolically distinct and resemble those in autoimmune disease. A) Bar plots of the expression of marker genes specific for EBV reprogrammed B cells in autoimmune disease in EBV^+^ and EBV^-^ B cells in patients with COVID-19. Genes are grouped by common function when possible (e.g. antigen presentation). B) Mountain plot of the enrichment of EBV reprogrammed B cell gene sets from Younis et al.^16^ for memory B cells (CD27^+^CD21^low^) and across all B cells. Normalized enrichment scores (NES) and FDRs are annotated on the plot. C) Venn diagram of the overlap between upregulated genes in EBV^+^ naïve B cells (red, upper left), memory B cells (blue, upper right), and plasmablasts (purple, lower). Representative genes upregulated in EBV^+^ B cells from all three cell states are annotated on the left. Representative genes upregulated in EBV^+^ memory B cells are annotated on the right. P-value was calculated using the Chi-Squared test. D) Contour plot of the distribution of EBV^+^ memory B cells on a UMAP of single memory B cell transcriptomes. Memory B cell states are outlined on the UMAP. E) Diagram of metabolic flux reactions related to glycolysis. Reactions (depicted by arrows) are annotated by their activity in EBV^+^ minus EBV^-^ B cells (positive values indicate increased activity in EBV^+^ B cells). F) Scatter plot of the difference in the average level of plasma long-chain carnitines (left) or dicarboxylic acids (right) in patients with high (>% reported on the X-axis) and low (<25^th^ percentile) proportions of EBV^+^ B cells (Y-axis). EBV^+^ high patients were filtered for progressively greater proportions of EBV^+^ B cells (X-axis); for example, X=N means we compared EBV low patients against EBV high patients who had at least N% of their B cells as EBV^+^. Differences that are significant (p<0.05) are indicated by a solid black outline and a dark blue color. Bar height represents arithmetic mean, and error bars represent standard error. P-values were calculated using the Mann-Whitney U test.

### EBV metabolically reprograms memory B cells in COVID-19

The enrichment of public EBV^+^ memory B cell gene signatures in the EBV^+^ B cells identified in this study (**Fig. 3B**) aligns with the known preference of EBV to remain latent in memory B cells. Consistently, in this study we found EBV^+^ memory B cells to present with the greatest upregulation of defining EBV reprogramming genes (e.g. *HLA-DRA*, *HLA-DRB1* encoding for antigen presentation machinery, **Fig. 3C**). Notably, we also observed significant overlap (p<0.001, by permutation) in the genes upregulated in EBV^+^ B cells across B cell states (**Table S11**). These genes were enriched for glycolytic and immunomodulatory functions (e.g. *ENO1*^49^ and *OAS1*^50^) and may suggest the existence of a core EBV reprogramming signature.

We then performed a targeted analysis of memory B cells, given their importance in EBV pathology. EBV^+^ memory B cells presented with a marked shift in phenotype upregulating genes related to B cell activation (e.g. *BTK*^44^*, NFKB1*^51^) and costimulation (e.g. *CD80, CD86*^52^) (**Figs. 3D and S4A-B**). The increased metabolic demands of cell activation, and subsequent proliferation, aligns with the upregulation of energy production pathways we observed in Figs. 2B and S2D. Together, this led us to hypothesize that EBV^+^ B cells may also undergo metabolic reprogramming, in the context of COVID-19. To address this, we performed metabolic flux analysis and found EBV^+^ memory B cells to significantly upregulate flux through the glycolysis pathway and decrease fatty acid breakdown (i.e. beta-oxidation) (**Fig. 3E, Table S12**).

With sufficient dysregulation, cellular metabolic reprogramming can lead to shifts in plasma metabolite levels. We assessed for such dysregulation in patients with increased proportions of EBV^+^ B cells and were indeed able to find sets of metabolites whose levels during convalescence (2-3 months after infection) significantly correlated with patient EBV^+^ B cell percentages (**Fig. S4C, Table S13**). The enrichment of these metabolites for dicarboxylic acids and carnitines, especially those of longer lengths, is consistent with our observation of reduced beta-oxidation in EBV^+^ memory B cells (**Fig. S4D-E**); these metabolites accumulate when beta-oxidation is impaired^53,54^. We confirmed this enrichment by finding long-chain carnitine and dicarboxylic metabolite modules to be significantly more abundant in patients with increased proportions of EBV^+^ B cells (**Figs. 3F and S4F**). These two metabolic module correlations suggest that the scRNAseq assessment of EBV-reprogrammed B cells is at least semi-quantitative. Notably, these metabolic modules are only mildly correlated with each other (shared variance r^2^=0.07 amongst patients with COVID-19), consistent with established biology that they report on distinct enzymatic routes. Thus, we find EBV^+^ memory B cells to undergo metabolic reprogramming at the cellular level, upregulated glycolysis programs and impaired beta-oxidation, that is reflected by global metabolite changes in plasma.

### EBV reprogrammed B cells provide stimulatory input to B and CD4^+^ T cells

The upregulation of costimulatory molecules in EBV^+^ B cells is reminiscent of reported mechanisms in autoimmune disease whereby EBV^+^ B cells activate autoreactive B and CD4^+^ T cells. This activation can lead to autoimmune pathology through, for example, the production of autoantibodies^16,55,56^. In conjunction with the shared clinical presentations of long COVID and autoimmune disease^57–60^, these data raise the possibility of shared molecular mechanisms between these conditions. To investigate this possibility, we queried for cell-cell interactions between EBV^+^ B cells and bystander EBV^-^ B and CD4^+^ T cells in patients with COVID-19 (**Table S14**). Strikingly, we observed EBV^+^ B cells to be significantly enriched for ligand-receptor interactions that facilitate B cell survival and activation (e.g. *BAFF:BAFFR*, *BAFF:TACI*)^61–65^ and CD4^+^ T cell co-stimulation (e.g. *CD80:CD28*, *CD86:CD28*^66,67^, *CD70:CD27*^68–70)^ (**Figs. 4A-B and S5**). In line with the role of this mechanism in autoimmune disease, we found patients with increased EBV^+^ B cell percentages to present with increased serum levels of autoantibodies (**Fig. 4C**) and increased symptomology 2-3 months after acute disease resolution (**Fig. S6**). Thus, in the context of COVID-19, EBV^+^ B cells appear to facilitate B and CD4^+^ T cell expansion through the expression of pro-survival and stimulatory ligands; this activation may facilitate the production of autoantibodies that can contribute to, certain, long COVID pathology (**Fig. 4D**)^71^.

**Fig. 4:**
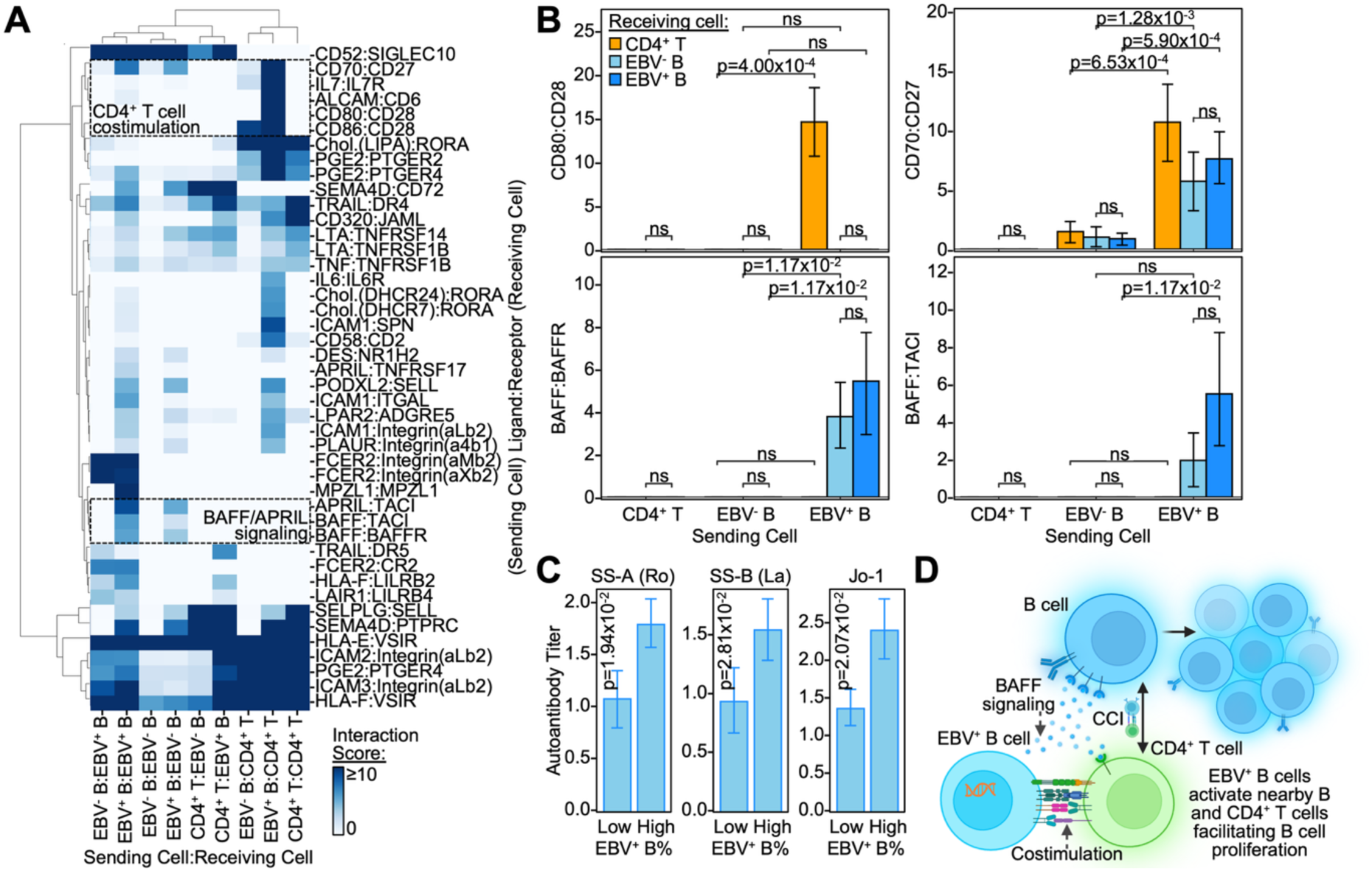
EBV reprogrammed B cells provide stimulatory signals to EBV^-^ B cells and CD4^+^ <u>T cells through cell-cell interactions.</u> A) Heatmap of ligand-receptor (rows) scores for pairwise interactions between EBV^+^ B cells, EBV^-^ B cells, and CD4^+^ T cells (columns). Cell color represents interaction score magnitude (see legend on the lower right). P-values that are non-significant (p≥0.05) are indicated with an annotation of “ns”. B) Bar plots of ligand-receptor interaction scores (Y-axis) between “sender” cells (X-axis) and “receiver” cells (represented by different colored bars). C) Bar plots of the serum titers of autoantibodies (Y-axis) in patients with high (>75^th^ percentile) and low (<25^th^ percentile) proportions of EBV^+^ B cells. D) Diagram of the cell-cell interactions identified between “sending” EBV^+^ B cells and “receiving” EBV^-^ B cells and CD4^+^ T cells. Bar height represents arithmetic mean, and error bars represent standard error. P-values were calculated using the Mann-Whitney U test.

### COVID-19 EBV^+^ B cell signature is enriched in patients with autoimmune disease

The molecular similarities between the EBV^+^ B cells in this work and EBV^+^ B cells in autoimmune disease^6,16,17,72^ suggest the existence of a core EBV reprogramming module across conditions (**Fig. S7A**). To investigate this, we interrogated for similarities between EBV^+^ B cells in COVID and in SLE, an autoimmune condition that shares some clinical features with long COVID^57–60^. We found the EBV^+^ B cell signature in COVID to significantly positively correlate with the expression of genes implicated in SLE (RNA, transcriptomic) and for patient EBV^+^ B cell percentages to significantly positively correlate with their polygenic SLE risk (DNA, genomic) (**Fig. 5A**). We confirmed these findings by showing our EBV^+^ B cell signatures in the COVID cohort to be significantly enriched in a public cohort of patients with SLE^73^ (**Figs. 5B-C and S7B-C, Table S15**). As EBV reprogramming is also implicated in MS, we repeated this analysis in a public cohort of patients with MS^74^ and similarly found enrichment of our EBV^+^ B cell signatures from the COVID cohort (**Fig. 5D**).

**Fig. 5:**
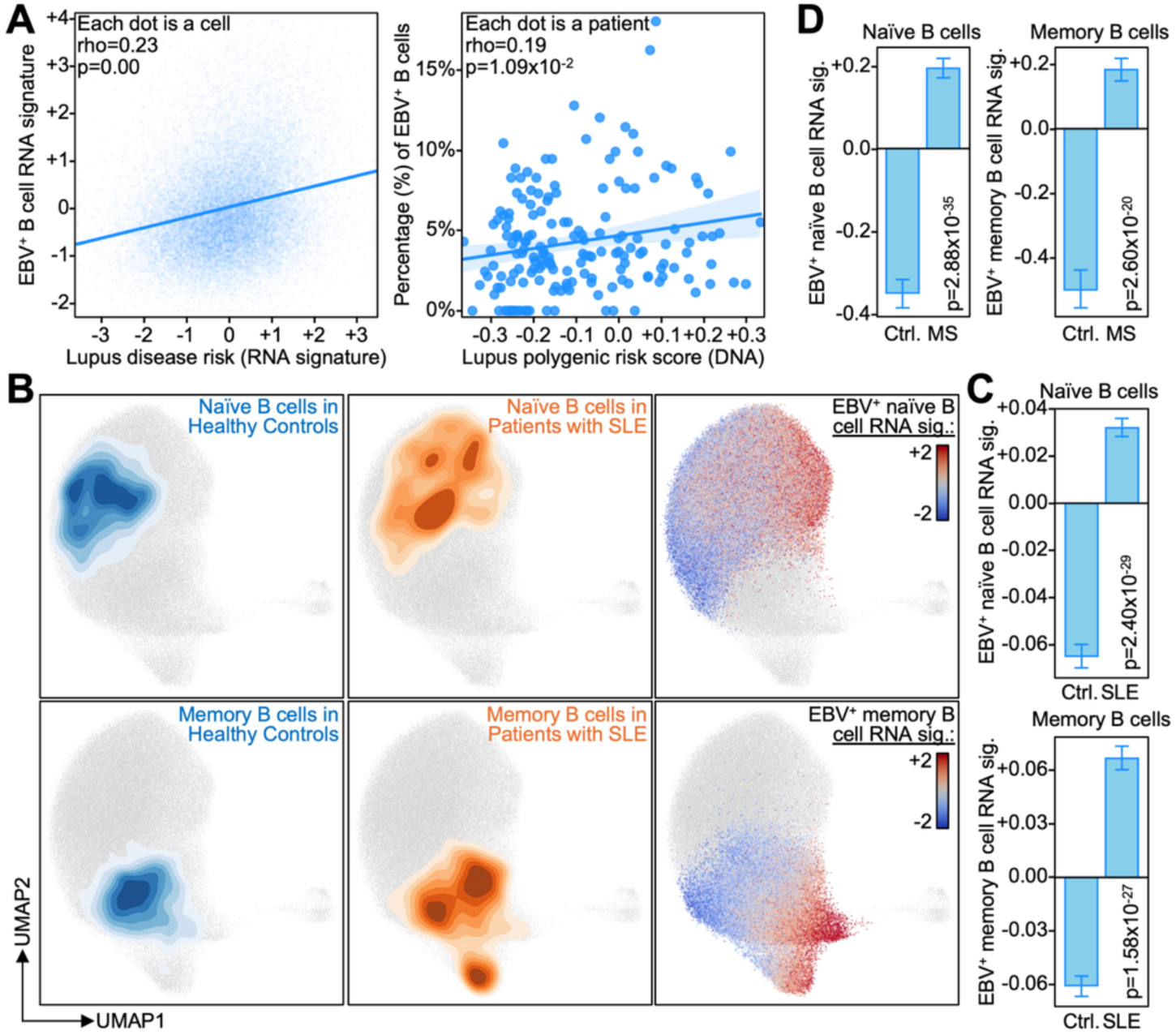
EBV reprogramming gene signatures in COVID are enriched in autoimmune disease. A) Left: Fitted line plot between single B cell expression of SLE disease relevance score (X-axis, from Zhang and Hou et al.^107^) and this work’s COVID-19 EBV reprogramming gene signature (Y-axis). Dots represent individual B cells. Right: Fitted line plot between a patient’s SLE polygenic risk (from DNA, whole genome sequencing) and the percentage of their B cell compartment that is EBV^+^ (Y-axis). Dots represent individual patients. Lines on the left and right represent fitted average and shaded areas represent 95% confidence interval. Pearson’s method was utilized to calculate correlation coefficients and p-values. B) Contour plot of the density of naïve (upper) and memory (lower) B cells from healthy controls (left) and patients with SLE (center) on a UMAP of single B cell transcriptomes. On the right, naïve and memory B cells are colored by their enrichment of this work’s B cell state specific EBV reprogramming signature in COVID (see legend on the right). C) Bar plot of the enrichment of this work’s B cell state specific EBV reprogramming signature in COVID (Y-axis) in naïve (upper) and memory (lower) B cells from patients with SLE compared to those from healthy controls (X-axis). D) Bar plot of the enrichment of this work’s B cell state specific EBV reprogramming signature in COVID (Y-axis) in naïve (left) and memory (right) B cells from patients with MS compared to those from healthy controls (X-axis). Bar height represents arithmetic mean, and error bars represent standard error. P-values were calculated using the Mann-Whitney U test.

## Discussion

In this work, we have identified EBV^+^ B cells in patients with COVID-19, even 2-3 months after acute disease resolution, and revealed how they are transcriptionally and metabolically reprogrammed to facilitate not only their own activation but also the activation of nearby EBV^-^ B and CD4^+^ T cells through potent stimulatory signals (**Fig. 6**). To this end, we have four fundamental observations. First, the identification of a non-zero (non-amplified) transcript signal within a B cell that is assignable to the EBV EBNA2 gene is sufficient to identify that cell as EBV-reprogrammed. Second, that reprogramming, investigated here within the context of COVID-19, strongly resembles EBV reprogrammed B cells in autoimmune disease and is indeed associated with increased autoantibody titers and long COVID symptomology (e.g. difficulty concentrating). Third, the scRNAseq-assessed level of B cell reprogramming is at least semiquantitative; the impact in patients with a higher % of EBV transformed B cells is measurably greater. Finally, reprogrammed B cells can exert significant influence over the global plasma environment of an individual.

**Fig. 6:**
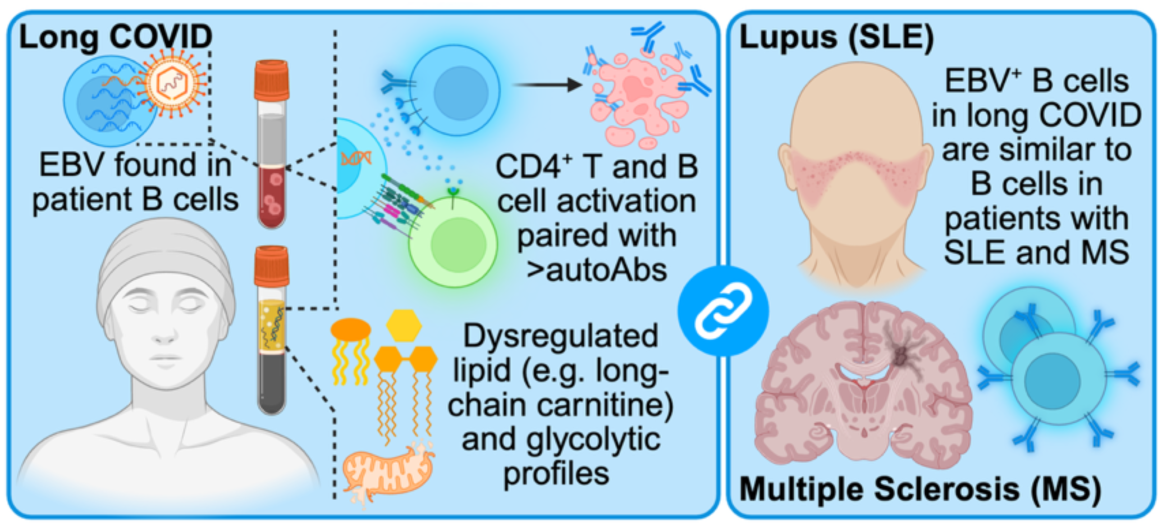
EBV reprogramming of B cells in COVID-19 and potential similarities to autoimmune disease. Diagram of the molecular, cellular, and plasma level characteristics of EBV reprogramming of B cells in patients with COVID-19 and similarities across these multi-omics to EBV reprogramming observed in autoimmune diseases.

Our finding that EBV^+^ B cells signatures in COVID-19 are similar to those observed in autoimmune disease, at the transcriptomic and epigenomic level (**Figs. 2C-E, S2B-D, 3A-B, and S3**), suggests the existence of a core EBV reprogramming gene module shared across diseases. This hypothesis is supported by the enrichment of certain transcriptional programs observed in this work and autoimmune disease in EBV-driven malignancies and lymphoproliferative disorders (e.g. MYC and NFkB signaling, apoptosis inhibition)^11,75^. This core module may be refined and leveraged by future works to 1) identify additional conditions where EBV reprogramming plays a role and 2) identify potential therapeutic targets expressed across EBV-associated pathologies. Further, the interrogation of gene expression changes in EBV^+^ B cells, by this work and others, will better our understanding of genes recently implicated as increasing risk EBV infection, and subsequent elevated risk of autoimmune disease and malignancies^76,77^. For example, decreased expression of *FOXO1* in EBV^+^ B cells after COVID-19 may align with the known repression of that gene by EBV LMP1/2 proteins (**Fig. S7D**)^78^. EBV^+^ B cell upregulation of *NFKB1* and *TNFRSF13B* proposes these genes as positive regulators of EBV persistence, potentially through their facilitation of B cell survival signaling^62,63^. As EBV^+^ B cells are more broadly identified and characterized in future studies, the cross-disease sharing and disease-specific roles of these genes will become ever more clear.

We reveal EBV^+^ B cell signatures in COVID-19 as similar to those in autoimmune disease at the transcriptomic level and in terms of their cell-cell communication profiles (**Figs. 3A-B and S5**). Recent works on SLE and MS demonstrated EBV^+^ B cells as drivers of autoimmune pathology by facilitating the expansion of both themselves and autoreactive double negative 2 (DN2, IgD^-^CD27^-^) and memory B cells into autoantibody secreting plasmablasts, secondary to stimulatory ligand-receptor interactions^16,36,79,80^. Here, we find these same interactions to be present between EBV^+^ B cells and EBV^-^ B and CD4^+^ T cells in patients with COVID-19. These interactions are an attractive explanation for the increased autoantibody levels observed in patients with COVID-19 with increased EBV^+^ B cell percentages (**Fig. 3C**). This explanation is supported by evidence of COVID-19 DN2 cells as potent secretors of autoantibodies^30^ and the observed persistence of EBV^+^ B cells during convalescence (2-3 months after acute disease resolution) when long COVID symptoms are present. Should DN2 B cells be stimulated by EBV^+^ B cells in patients with COVID-19, as suggested by the EBV^+^ and EBV^-^ B cell interactions reported here, then there would exist a shared mechanism of autoantibody production across SLE, MS, and long COVID. This mechanism is supported by a notable similarity between the susceptibility of COVID-19 DN2 B cells to differentiate into autoantibody secreting plasmablasts and increased production of autoantibody secreting plasmablasts in SLE^81^. Future efforts analyzing these disease states in parallel may more formally elucidate the presence or absence of this mechanism.

We also find elevated fractions of EBV^+^ B cells to be associated with increased levels of both long-chain carnitine and dicarboxylic acid levels (**Figs. 3F and S4C-F**). These metabolite levels may be associated with the biology of EBV-driven B cell reprogramming, that includes increased glycolysis and reduced beta-oxidation. Impaired beta-oxidation, often secondary to mitochondrial stress, can lead to elevated levels of carnitines and dicarboxylic acids^53,54,82,83^. This accumulation can occur due to a build-up of reactants (e.g. fatty acid associated carnitines) and shunting towards omega-oxidation, which produces dicarboxylic acids as a product. Future metabolically-focused studies may explore the exact processes that underpin this metabolic stress and provide deeper characterizations of the alternate pathways (e.g. omega-oxidation) that are activated as a result. The observation of serum metabolite changes as a reflection of cellular dysfunction is not without precedent. Certain inborn errors of metabolism (e.g. medium-chain acyl-CoA dehydrogenase deficiency) that lead to cellular dysfunction can be diagnosed through plasma metabolite profiles, such as the distribution of carnitines^84,85^. It will be important for future studies to validate these findings in larger cohorts with more comprehensive metabolic panels. Positive reports from these validation studies would support the use of serum metabolite levels as a biomarker for certain types of long COVID and EBV-associated disease and propose the treatments used for similar congenital metabolic disorders as potential therapeutics for these conditions.

Together this work synergizes with existing literature to propose a potentially addressable mechanism of long COVID: EBV^+^ B cell driven production of autoantibodies, recently reported to be causative agents of long COVID symptoms^71,86^, and concomitant immune cell and metabolic dysregulation, potentially secondary to interactions with EBV^+^ B cells. Together, these data suggest the clearance of EBV^+^ B cells, or disruption of their signaling, as a potential therapy for long COVID and, given potentially shared molecular mechanisms and clinical features, autoimmune disease.

### Limitations of the Study

This study is limited by the number of identified EBV^+^ B cells. Our approach leveraged scRNAseq data that did not undergo amplification for EBV reads (via EBV-specific primers). This has advantages in terms of extracting quantitative relationships, but only cells with markedly high expression of EBV transcripts could be assigned as EBV^+^. Further, due to the limited number of EBV^+^ B cells in healthy donors, we were restrained in our ability to comment on the EBV reprogramming differences between healthy donors and patients with COVID-19. This may be elaborated in future works with larger cohorts of healthy donors where deep B cell sequencing may identify these rare EBV^+^ B cells. Future studies with EBV-specific primers may reveal rare EBV^+^ B cells in patients with COVID-19 that could not be observed in this work. While we validated our findings in two cohorts of patients with COVID-19, validation of these findings in additional cohorts can help assess the generalizability of our findings. Similarly, the autoimmune disease comparisons in this work were largely with SLE and MS, though additional autoimmune diseases were compared against via disease risk scores in **Fig. S7A**. Future works may seek to perform comparisons with additional autoimmune diseases as applicable data becomes available. Last, this study focuses on transcriptomic differences between EBV^+^ and EBV^-^ B cells, with metabolic differences inferred from said transcriptomic data. Emerging technologies that can capture multiple modalities with deep coverage will allow for more mechanistic understandings of EBV reprogramming of B cells.

## Methods

### Identification of EBV^+^ cells from scRNAseq data

EBV read detection was performed on two public cohorts of patients with COVID-19 and healthy controls (INCOV, HAARVI)^25^. EBV reads were identified using a previously published computational framework^16^. In brief, binary alignment map (BAM) files were retrieved from standard 10x Genomics cellranger output (“possorted_genome_bam.bam”) and filtered for unmapped reads (i.e. those not mapped to the human genome) via SAMtools (v1.23.1)^87^ with “samtools view -b -f 4 INPUT_FILE > OUTPUT_FILE”. Non-human reads were then mapped against the EBV genome (NCBI reference sequence: NC_007605.1). EBV gene annotations, in gene transfer format (GTF), were refined as per Younis et al.^16^ and Grande and Gerhard et al.^88^. Read mapping was performed using Spliced Transcripts Alignment to a Reference (STAR, v2.7.10a)^89^ with parameter values set to those defined by Younis et al.^16^; cell barcode whitelists were downloaded from 10x Genomics (“737K-august-2016.txt” was utilized for the 10x chemistry utilized in this work). Cell barcodes were gathered from reads that aligned to the EBV genome and visualized via the Integrative Genomics Viewer (IGV, v2.11.9)^90^ after deepTools (v3.5.6)^91^ processing.

### Assignment of B cell states

Single B cell transcriptome UMAP positions and unbiased clustering assignments (calculated via the leiden^92^ algorithm) were previously reported^25^. B cell marker genes were utilized to annotate unbiased clusters into B cell states: *FCER2* (encoding CD23) and *IGHD* (encoding IgD) for naïve B cells, *IGHM* (encoding IgM) for unswitched memory B cells, *MS4A1* (encoding CD20) for non-plasmablast B cells, *ZEB2* and *FCRL5* for atypical memory (double negative 2) B cells, and *XBP1* and *CD38* for plasmablasts. B cell states that were heterogenous (composed of multiple unbiased clusters) were separated with a numerical suffix (e.g. Naïve I, Naïve II, Naïve III).

### Differential gene expression analysis

Differentially expressed genes were identified through pairwise comparisons between the expression of each gene in group A (e.g. EBV^+^ B cells) and group B (e.g. EBV^-^ B cells). Expression values were normalized by library size (counts per million plus a pseudocount for one, as previously reported^25^) and a log-transformation (natural base). Normalized values are utilized for all statistical analyses, imputed (by MAGIC^93^) expression was only utilized for visualization purposes on single cell UMAPs. P-values were calculated using the Mann-Whitney U test and corrected for multiple hypothesis testing using the Benjamin-Hochberg method (also called False Discovery Rate or FDR).

### Gene set enrichment analysis

Differentially expressed genes underwent gene set enrichment analysis as implemented by EnrichR^94^ for the enrichment of gene sets from discrete gene lists (top 500 genes unless otherwise specified) or GSEApy (v1.2.1)^95^ for the enrichment of gene sets in continuously ranked gene lists (as ranked by Mann-Whitney U statistic unless otherwise specified). Transcription factor program enrichment analysis was performed using ChEA^96^. Pathway enrichment analysis was performed using KEGG^97^ and GO Biological Process^98^. Kinase enrichment analysis was performed using ARCHS4^99^. Disease association enrichment analysis was performed using Orphanet^100^ and GWAS Catalog^101^. These databases are natively provided by EnrichrR. The SLE EBV^+^ memory B cell (CD27^+^CD21^low^) gene set was manually curated from Fig. 3B by Younis et al.^16^. The SLE EBV^+^ all B cell gene set was also curated from Younis et al.^16^ and defined as the union of the gene sets in differentially upregulated pathways in EBV^+^ B cells across B cell states. Continuous gene set enrichment scores (EBV^+^ B cell, EBV^+^ naïve B cell, and EBV^+^ memory B cell signatures after COVID) were calculated using Scanpy’s (v1.10.3)^102^ “scanpy.tl.score_genes” function with the top 500 upregulated genes used as input (e.g. top 500 upregulated genes in EBV^+^ memory B cells, compared to EBV^-^ memory B cells, for the EBV^+^ memory B cell signature after COVID). Gene signatures were normalized to standardized Z-scores prior to downstream analyses.

### Chromatin openness and transcription factor binding analyses

Single cell multiome (simultaneously measured RNA- and ATAC-seq information from the same single cell; i.e. sc-multiome) data of B cells from patients with COVID-19 was acquired from our recently reported work^30^. EBV^+^ B cells were identified from the transcriptome portion of the sc-multiome data as per the section “Identification of EBV+ cells from scRNAseq data”. Epigenetic (ATAC-seq) reads from identified EBV^+^ and EBV^-^ B cells were isolated using the subset-bam (v1.1.0) program from 10x Genomics. Subsetted BAM files were then processed into BigWig files via deepTools (v3.5.6)^91^ using reads-per-kilobase-million normalization. Peaks identified by the cellranger-arc (v2.0.2) package from 10x Genomics were utilized for differential chromatin openness analysis whereby peaks were identified as being significantly open (p<0.05) via the Mann Whitney U test. Differentially open peaks were then submitted to MEME (v5.5.9)^103^ with the motif position-weighted-matrices in MEME format for EBF1 (MA0154.2) and RBPJ (MA1116.1) acquired from the JASPAR database^104^.

### Metabolic flux and metabolomics analysis

Single cell metabolic flux analysis was performed using scFEA^105^. scFEA provides calculated flux values for individual reactions (e.g. conversion of fatty acids to acetyl-CoAs via beta-oxidation). As scFEA requires imputed expression values, these were calculated using MAGIC^93^, as recommended by the package. Metabolomic analyses were performed on previously reported Metabolon data^25^. We identified metabolites associated with increased proportions of EBV^+^ B cells in patient B cell compartments via correlation analysis (Pearson’s method). This was performed between a patient’s maximum EBV^+^ B cell percentages cross timepoints and timepoint specific metabolite levels in plasma. The EBV^+^ B cell percentages described here were utilized for all patient level comparisons (e.g. with maximum autoantibody levels in patient serum).

### Cell-cell communication analysis

Cell-cell communication analysis was performed via CellphoneDB (v5)^106^ using their method “cpdb_statistical_analysis_method”. Known ligand-receptor pairs are natively provided by CellphoneDB. Individual blood draws were treated as separate microenvironments, as cell-cell interactions cannot occur between patients nor transcend timepoints. Default method parameters were utilized and no subsampling was performed. CellphoneDB calculated interaction scores were utilized to identify ligand-receptor interactions differentially prevalent in cell-cell interactions between EBV^+^ B cells and other cell types (EBV^+^ B cells, EBV^-^ B cells, and CD4^+^ T cells).

### Association of EBV^+^ B cell percentage with post-acute symptomology

Post-acute (2-3 months after acute disease resolution) symptom data was acquired from our previous work^25^. EBV^+^ B cell percentage data was computed per sample; samples with less than 50 B cells were not utilized for analyses as their lower coverage of sample B cells precludes accurate assessment of EBV^+^ B cell proportion. The Mann-Whitney U test was utilized to identify symptoms where there was a significant difference in EBV^+^ B cell proportion in patients with and without said symptom.

### Comparisons between EBV^+^ B cells in COVID and autoimmune disease

Comparisons to autoimmune disease were performed by quantifying the expression of polygenic disease risk scores, as calculated by scDRS (v1.0.2)^107^, in COVID-19 B cells. Single cell level comparisons were performed in continuous fashions, correlation (Pearson’s method) between expression of autoimmune disease risk scores and the COVID-19 EBV^+^ B cell gene signature reported in this work, and discrete fashions, calculating for statistical differences in the expression of autoimmune disease risk scores in EBV^+^ and EBV^-^ B cells after COVID with p-values calculated by the Mann-Whitney U test. Patient level comparisons were performed using previously reported polygenic risk scores^30^ and patient EBV^+^ B cell percentages (as defined in the “Metabolic flux and metabolomics analysis” Methods section: the maximum proportion of a patient’s B cell compartment that was EBV^+^ across timepoints).

Public scRNAseq data from patients with SLE and healthy controls^108^ was downloaded from CELLxGENE^109^ as an H5AD object. Author annotated B cells (denoted by “B” and “PB” labels) were subsetted for and subject to batch-corrected k-nearest-neighbor graph calculation (via bbkNN^110^ with batch set to “Processing_Cohort”), two-dimensional UMAP^111^ projection, and unbiased clustering (via the leiden^112^ algorithm). B cell marker genes (e.g. *TCL1A*, *FCER2*, *CD27*, *FCRL5*) were utilized to define unbiased clusters representing naïve B cells (clusters 0, 1, 2, 4, and 6) and memory B cells (clusters 3, 5, and 7). Public scRNAseq data from patients with MS^74^ and controls with idiopathic intracranial hypertension was downloaded from GEO^113^ (GSE138266) and subject to typical scRNAseq processing: filtering for high-quality cells (≥1000 unique molecular identifiers per cell, ≥500 unique genes per cell, <10% mitochondrial reads per cell) and removal of doublets (via Scrublet^114^) with a maximum doublet score set to 0.25. B cells were identified as unbiased clusters (calculated via the leiden^112^ algorithm) with upregulation of B cell marker genes *CD79A* and *CD79B*. Unbiased clustering was performed on a k-nearest-neighbors graph calculated from principal component analysis (PCA) dimensions; PCA was performed on highly variable genes as defined by “scanpy.pp.highly_variable_genes” (which was called with default parameters). EBV^+^ naïve and memory B cell gene signatures after COVID were calculated in the same manner as described in the “Gene set enrichment analysis” Methods section.

## Quantification and statistical analysis

P-values were calculated as follows: for pairwise comparisons of continuous values the Mann-Whitney U test was utilized, for comparisons of the difference in means across multiple groups (greater than two) the Kruskal-Wallis test was utilized, and for overlap between multiple sets (greater than two) the Chi-squared test was utilized. P-values were corrected for multiple hypothesis testing via the Benjamin-Hochberg method (FDR) when multiple tests were performed. Correlations (and associated correlation coefficients and p-values) were calculated via Pearson’s method. All single-cell analyses were performed using Scanpy (v1.10.3)^102^. All patient level analyses were performed with filtering for samples with at least ten cells.

## Data Availability

All analyses were performed on publicly-available data^25,73,74,115^; relevant calculated values (e.g. EBV^+^ B cell assignment) are provided as Supplementary Tables.

## Code Availability

This paper does not report original code. All work leveraged previously reported pipelines or publicly-available packages. Any additional information required to reanalyze the data reported in this work is available from the Lead Contact upon reasonable request.

## Acknowledgements

We are thankful for kind and insightful discussions from Dr. Heather E. Bukiri, Dr. Bevra Hahn, Dr. Katie M. Campbell, and Serey Nouth. D.G.C. is supported by the UCLA-Caltech Medical Scientist Training Program and Paul and Daisy Soros Fellowship. Y.S. is supported by the Damon Runyon Quantitative Biology Fellowship from the Damon Runyon Cancer 602 Research Foundation (DRQ-13-22), the Mahan Fellowship at Herbold Computational Biology Program of Fred Hutchinson Cancer Research Center, the Translational Data Science Integrated Research Center New Collaboration Award and Integrated Research Center at Fred Hutchinson Cancer Research Center, Pilot Award of Immunotherapy Integrated Research Center, and in part through the NIH/NCI Cancer Center Support Grant P30 (CA015704). J.D.G. is supported by the Swedish Medical Foundation. J.R.H., J.D.G. and H.C. are supported by the NIH RECOVER Initiative (OT2HL161847-01), and the INCOV cohort was supported by the Biomedical Advanced Research and Development Authority (HHSO10201600031C), the Wilke Family Foundation, the Murdock Trust, and Gilead Sciences.

## Author Contributions

Conceptualization: D.G.C., Acquired Data: J.D.G., H.C., Data Curation: D.G.C., Y.S., J.R.H., Formal Analysis: D.G.C., J.R.H., Investigation: D.G.C., J.R.H., Visualization: D.G.C., Funding Acquisition: J.R.H., Resources: J.R.H., Writing - original draft: D.G.C., Writing - review & editing: D.G.C., Y.S., H.C., J.D.G., J.R.H., Project Administration: J.R.H., Supervision: Y.S., J.R.H.

## Supplementary Figures

**Fig. S1:**
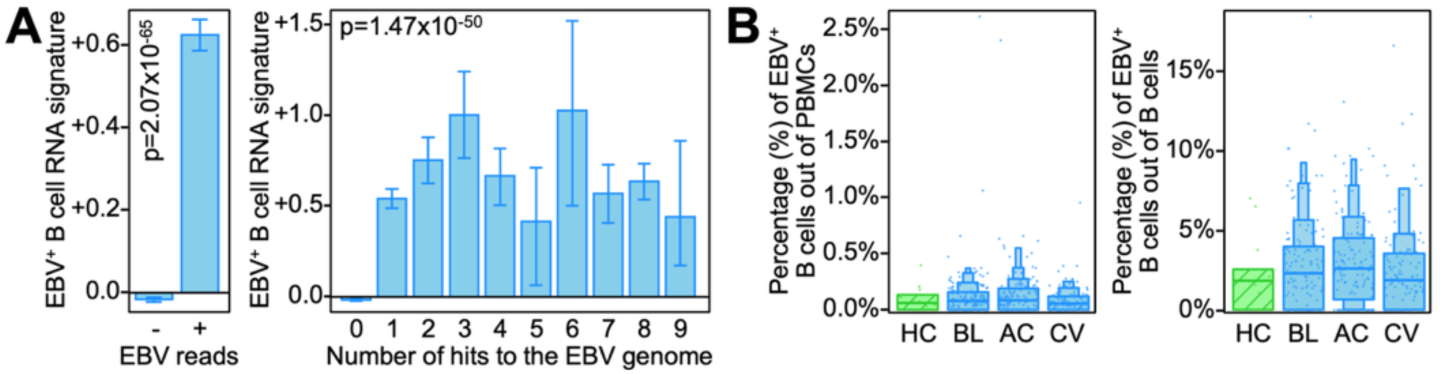
Computational EBV^+^ B cell assignment and validation. A) Bar plots of the enrichment of an EBV gene signature (derived from differentially expressed genes in EBV^+^ B cells) between EBV^-^ and EBV^+^ B cells (left) and EBV^-^ B cells and EBV^+^ B cells stratified by the number of times their EBV viral reads mapped to the EBV genome (right, e.g. 1=unique mapping, N=mapped N times). P-values were calculated using the Mann-Whitney U test (left) and Kruskal-Wallis test (right). B) Boxen plots of the percentage of a given sample (blood draw) with detectable EBV reads out of all PBMCs (left) or B cells (right). Samples are stratified by disease state: healthy donors (HD), baseline (BL, diagnosis of COVID-19), acute disease (AC, one week after diagnosis), and convalescence (CV, 2-3 months after acute disease). Boxen plots represent interquartile range (25 to 75 percentiles) with successive boxes representing progressively halved quantiles (e.g. 12.5/87.5, 6.25/93.75 percentiles). Dots represent individual samples.

**Fig. S2:**
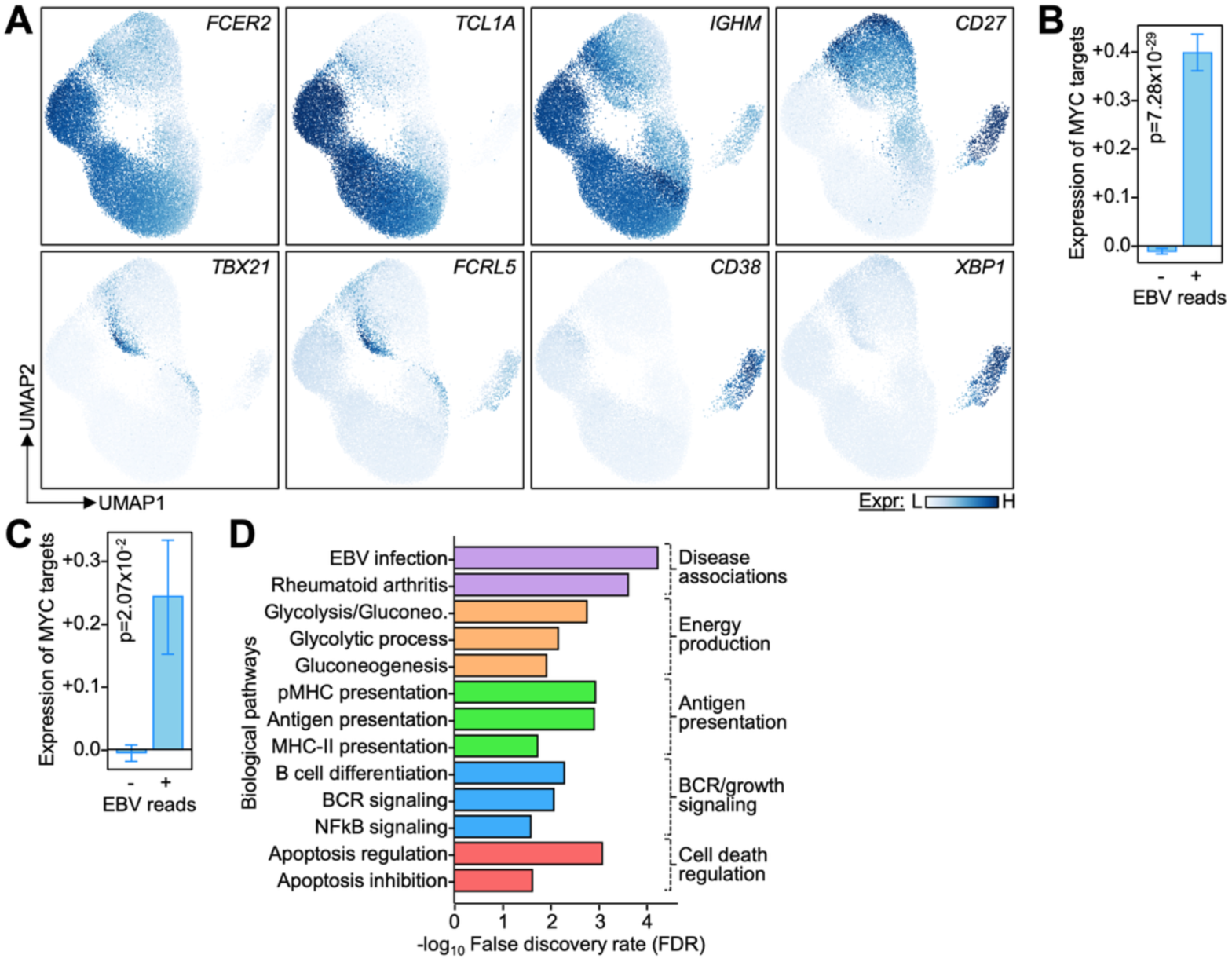
EBV transcriptionally reprograms B cells in patients with COVID-19. A) UMAP of single B cell transcriptomes colored by their expression of B cell marker genes (see legend on the right). B) Bar plot of the enrichment of the MYC transcription factor program (from the Hallmark collection of the MSigDB database) in EBV^+^ and EBV^-^ B cells. C) Bar plot of the enrichment of the MYC transcription factor program (from the Hallmark collection of the MSigDB database) in EBV^+^ and EBV^-^ B cells from the validation cohort (HAARVI). D) Bar plot of the pathways enriched in the differentially expressed genes in EBV+ B cells compared to EBV^-^ cells in the validation cohort (HAARVI, on the Y-axis) as quantified by -log10(FDR) (on the Y-axis). Color represents shared pathway categories (e.g. antigen presentation). Bar height represents arithmetic mean, and error bars represent standard error. P-values were calculated using the Mann-Whitney U test.

**Fig. S3:**
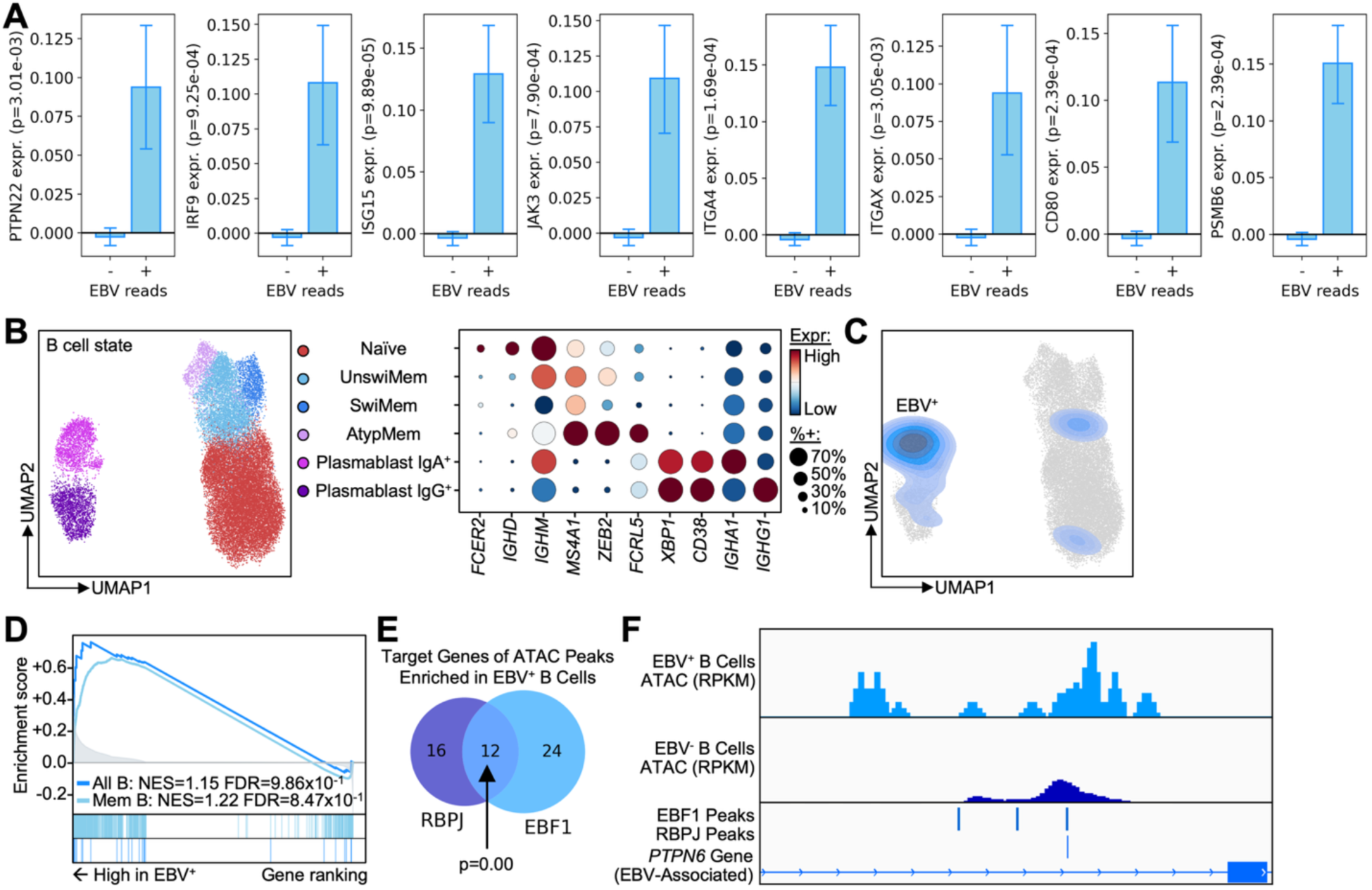
EBV^+^ B cells in patients with COVID have EBV reprogramming signatures in their <u>transcriptome and epigenome.</u> A) Bar plots of the expression of marker genes specific for EBV reprogrammed B cells in autoimmune disease in EBV^+^ and EBV^-^ B cells in patients with COVID-19. B) Left: Uniform manifold approximation projection (UMAP) of single B cell multiomes; transcriptome (used for embedding) and epigenome were simultaneously measured from the same single cell. Cells are colored by their cell state (see legend on the right); unswitched memory is abbreviated as UnswiMem, switched memory as SwiMem, and atypical memory as AtypMem. Right: Dot plot of the expression of B cell marker genes (columns) by cell state (rows). Dot size represents the percent of a given cell state that expresses said marker gene and color represents expression magnitude (see legend on the right). C) Contour plot of the distribution of EBV+ B cells on the UMAP from panel (B). D) Mountain plot of the enrichment of EBV reprogrammed B cell gene sets from Younis et al.^16^ for memory B cells (CD27^+^CD21^low^) and across all B cells. Normalized enrichment scores (NES) and FDRs are annotated on the plot. E) Venn diagram of the number of differentially open chromatin (“peaks”) in EBV^+^ B cells enriched for RBPJ and EBF1 transcription factor binding sites. P-value was calculated using the hypergeometric overlap test. F) Track plots of the pseudobulk chromatin landscape (ATAC-seq portion of single cell multiome from panel [B]) of EBV^+^ (upper, light blue) and EBV^-^ (lower, dark blue) B cells. Differentially open chromatin (“peaks”) significantly enriched for EBF1 and RBPJ binding sites are annotated on the plot. *PTPN6* gene body (derived from RefSeq) is annotated on the plot at the bottom.

**Fig. S4:**
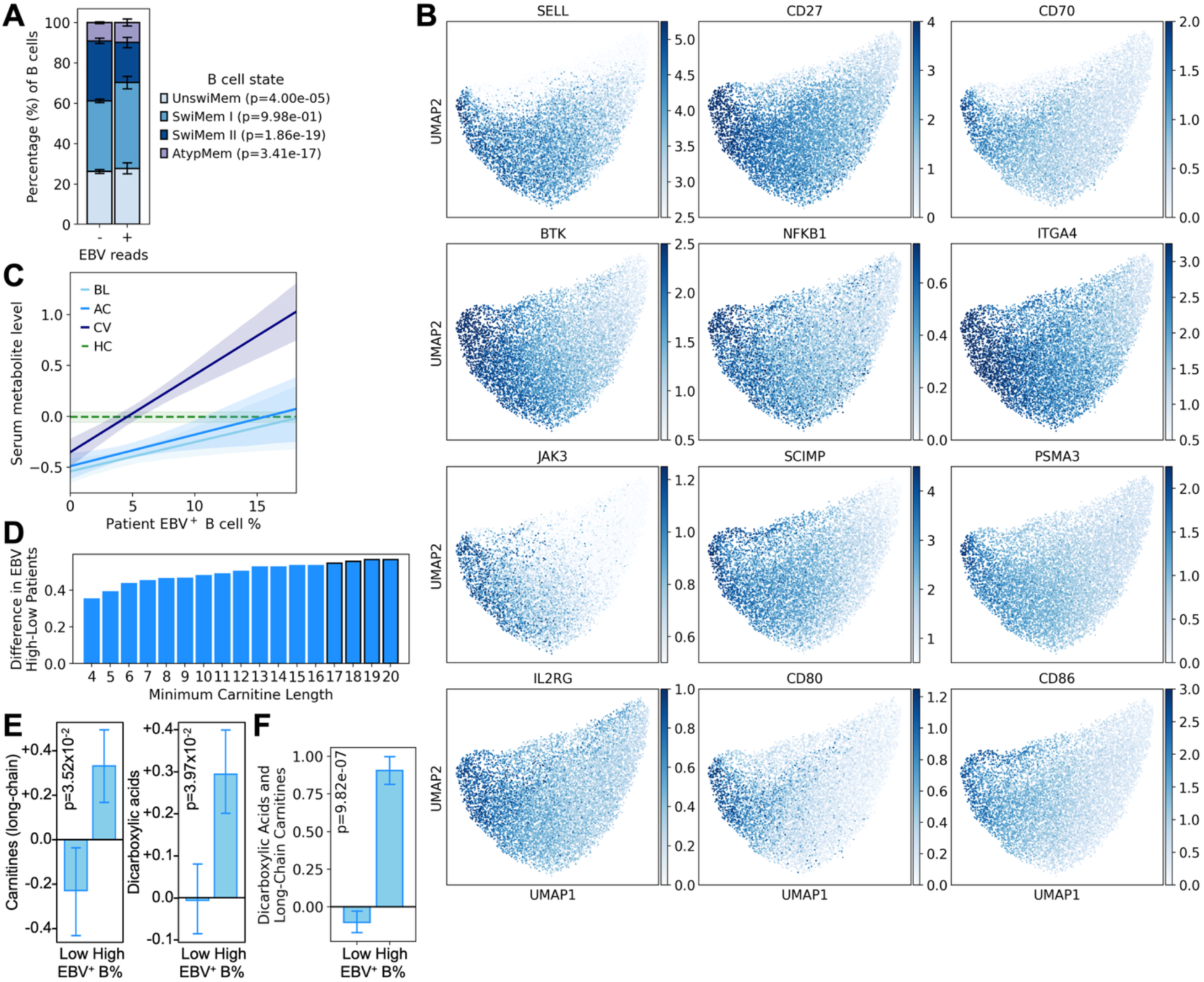
EBV^+^ B cells in patients with COVID-19 bear distinct transcriptomes and <u>metabolic states reminiscent of those in autoimmune disease.</u> G) Stacked bar plots of the distribution of memory B cell states amongst EBV^-^ and EBV^+^ memory B cells. Bars are colored by cell state (see legend on the right). H) UMAP of single memory B cell transcriptomes colored by their expression of memory B cell marker genes (see legend on the right). I) Fitted line plot between the percentage of a patient’s B cells that are EBV^+^ (X-axis) and level of selected metabolites in plasma (Y-axis) at diagnosis (BL, rho=0.18 p-value=1.70×10^-2^), during acute disease (AC, rho=0.20 p-value=9.08×10^-3^), and convalescence (CV, rho=0.50 p-value=1.33×10^-8^) (see legend on the upper left). Line represents fitted average and shaded area represents 95% confidence interval. Average (dashed green line) and 95% confidence interval (shaded green area) of said metabolites in healthy controls (HCs) are projected on the plot as a reference. Selected metabolites are those whose CV levels significantly positively correlate with patient EBV^+^ B cell percentages. J) Bar plot of the difference in the average level of plasma carnitines in patients with high (>75^th^ percentile) and low (<25^th^ percentile) proportions of EBV^+^ B cells (Y-axis). Carnitines are filtered for progressively longer lengths (X-axis); for example, X=N means carnitines with a length of at least N are utilized for Y-axis calculation. Differences that are significant (p<0.05) are indicated by a solid black outline. K) Bar plots of the average plasma levels of long-chain (LC, ≥18 carbons) carnitines and dicarboxylic acids in patients with high (>75^th^ percentile) and low (<25^th^ percentile) proportions of EBV^+^ B cells. L) Bar plot of the average plasma levels of a module of long-chain (LC, ≥18 carbons) carnitines and dicarboxylic acids in patients with high (>10%) and low (<25^th^ percentile) proportions of EBV^+^ B cells. Bar height represents arithmetic mean, and error bars represent standard error. P-values were calculated using the Mann-Whitney U test.

**Fig. S5:**
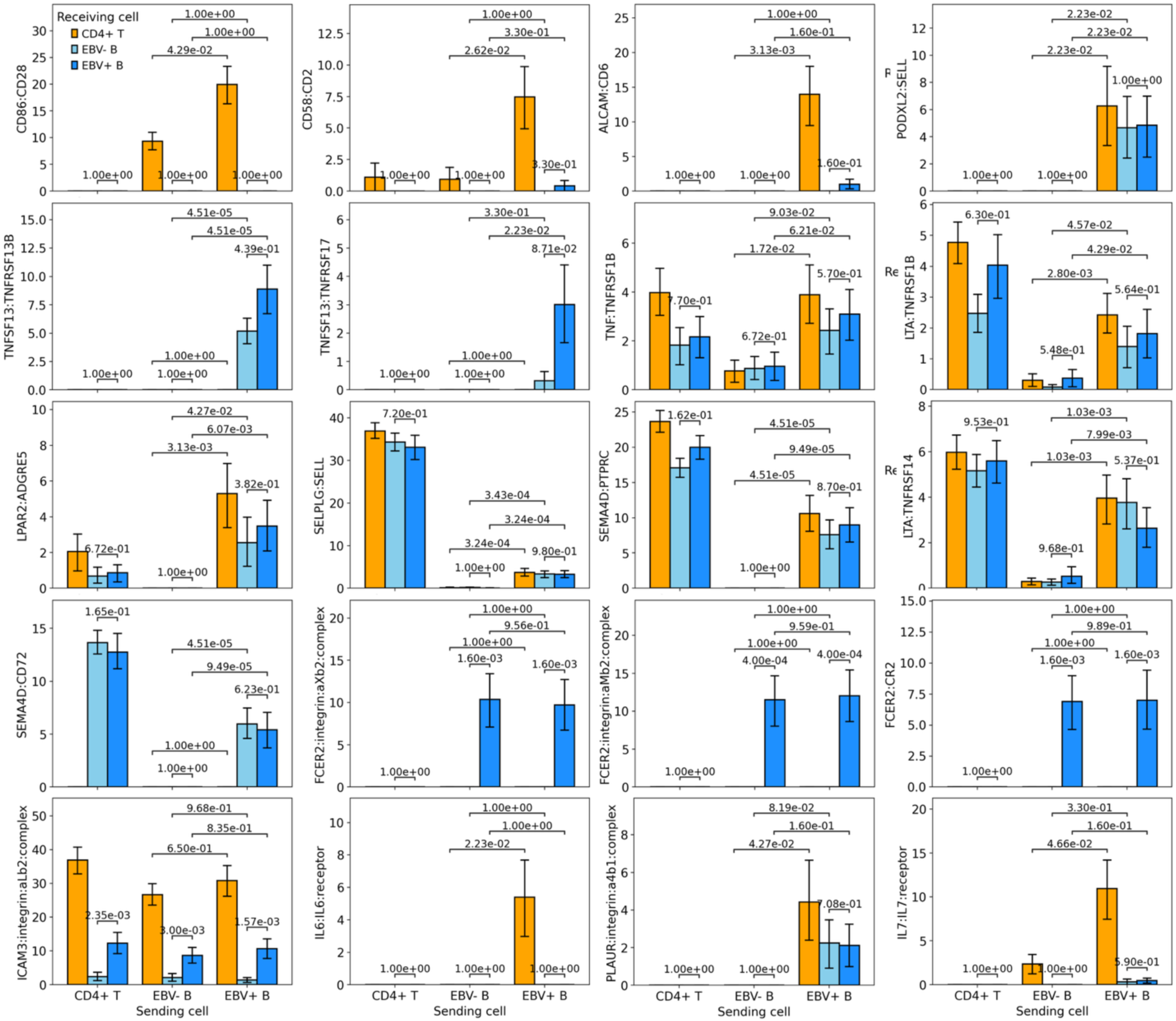
EBV^+^ B cells provide survival and costimulatory signals to EBV^-^ B cells and CD4^+^ <u>T cells.</u> Bar plots of ligand-receptor interaction scores (Y-axis) between “sender” cells (X-axis) and “receiver” cells (represented by different colored bars). Bar height represents arithmetic mean, and error bars represent standard error. P-values were calculated using the Mann-Whitney U test.

**Fig. S6:**
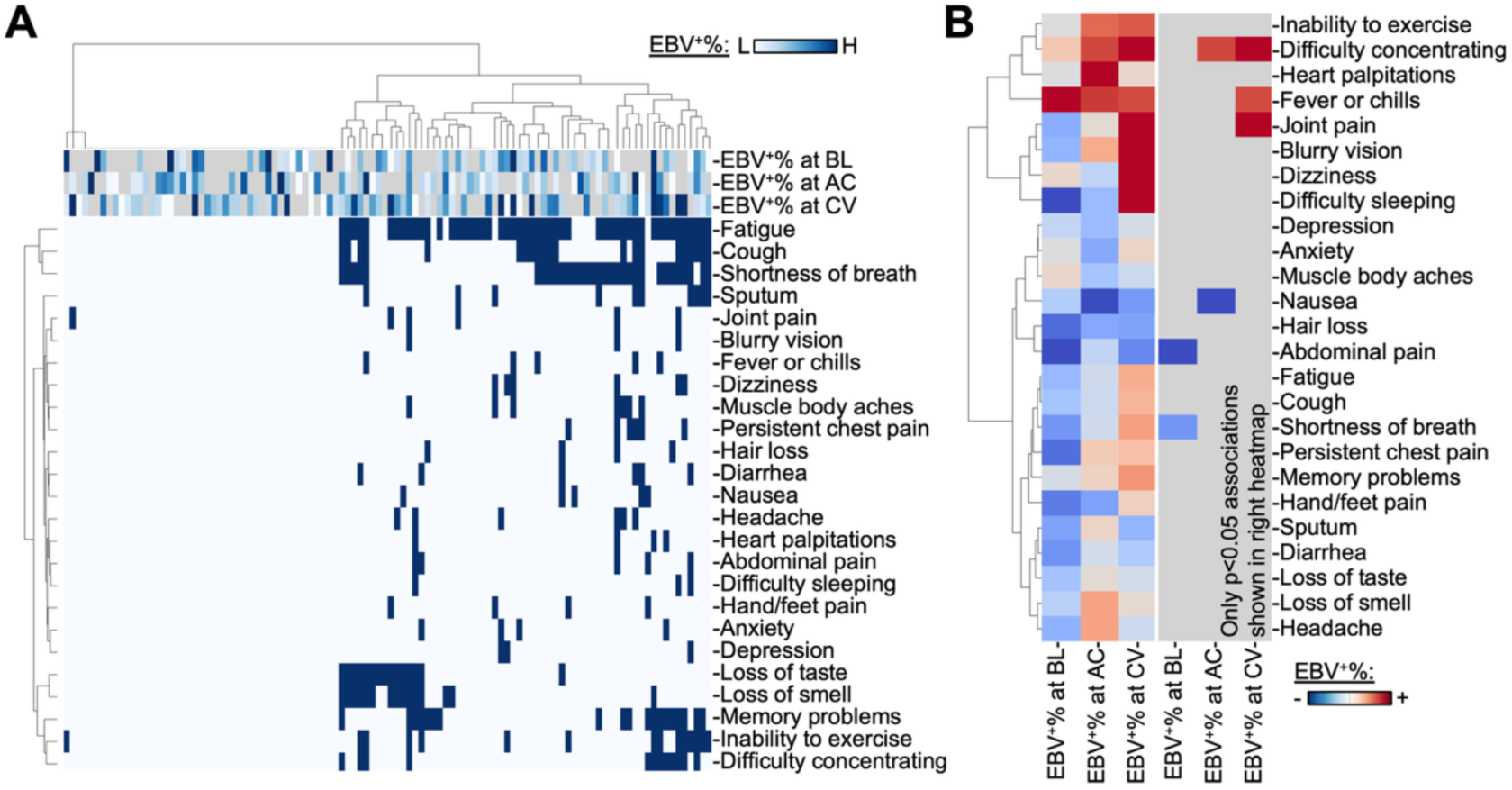
Patients with COVID-19 with higher proportions of EBV^+^ B cells had increased symptomology 2-3 months after acute disease resolution. A) Heatmap of the presence (dark blue) or absence (light blue) of each symptom (rows) for each patient (columns) 2-3 months after acute disease resolution (i.e. convalescence). Each sample’s EBV^+^ B cell percentage is annotated on the top (see color bar on the upper left); gray indicates insufficient number of B cells to calculate EBV^+^ B cell proportion. B) Heatmap of the average difference in EBV^+^ B cell percentage in patients with and without a given symptom (rows) at each timepoint (columns); BL is baseline at diagnosis, AC is acute disease, and CV is convalescence 2-3 months after acute disease resolution. Red color indicates an increased proportion of EBV^+^ B cells in patients with a given symptom, see color bar on the lower right. All differences are presented on the left, significant differences (p<0.05) are present on the right. P-value were calculated using the Mann-Whitney U test.

**Fig. S7:**
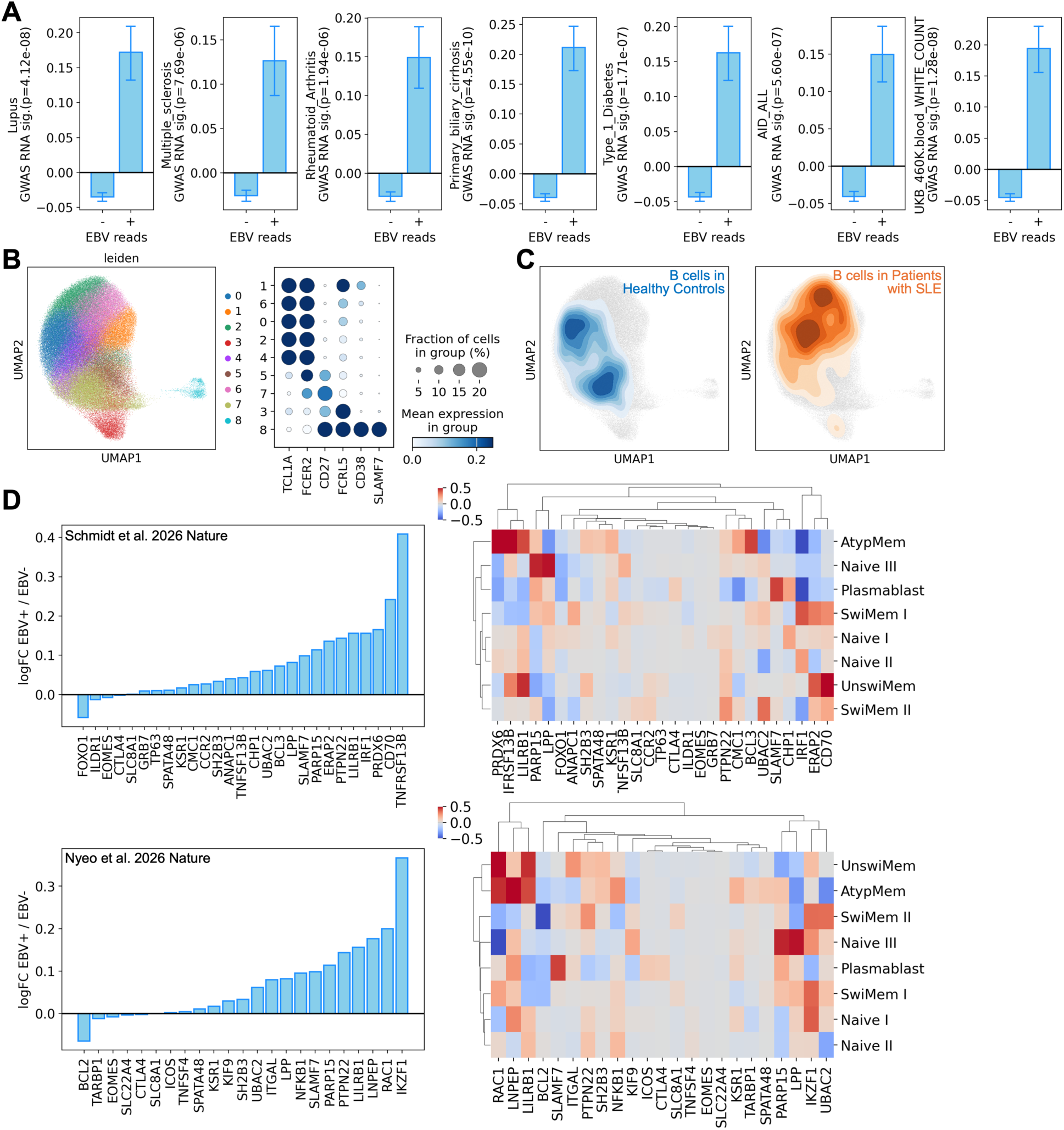
EBV^+^ B cells in COVID-19 are enriched for autoimmune disease features and genes implicated in EBV viremia by GWAS. A) Bar plots of the enrichment of autoimmune disease relevance scores (method by Zhang and Hou et al.^107^) in EBV^+^ and EBV^-^ B cells in patients with COVID-19. Bar height represents arithmetic mean, and error bars represent standard error. P-values were calculated using the Mann-Whitney U test. B) Left: UMAP of single B cell transcriptomes from patients with SLE and healthy controls. Cells are colored by their cell state (see legend on the right). Right: Dot plot of the expression of B cell marker genes (columns) by cell state (rows). Dot size represents the percent of a given cell state that expresses said marker gene and color represents expression magnitude (see legend on the right). C) Contour plot of the density of B cells from healthy controls (left) and patients with SLE (right) on the UMAP from panel (B). D) Left: Expression of genes (X-axis) implicated in EBV viremia in population scale studies by Schmidt et al.^76^ (upper) and Nyeo et al.^77^ (lower) in EBV^+^ minus EBV^-^ B cells; positive values indicate increased expression in EBV^+^ B cells. Right: Heatmap of the expression of genes (columns) implicated in EBV viremia from population scale studies by Schmidt et al.^76^ (upper) and Nyeo et al.^77^ (lower) in EBV^+^ minus EBV^-^ B cells across B cell states (rows); positive values indicate increased expression in EBV^+^ B cells (see legend on the upper left).

## Supplementary Tables

Table S1: EBV^+^ B cell assignment. scRNAseq annotation of B cells sample of origin and EBV positivity assignment.

Table S2: Differentially expressed genes in EBV^+^ B cells. Differential gene expression analysis between EBV^+^ and EBV^-^ B cells in the INCOV cohort.

Table S3: Transcription factor enrichment in EBV^+^ B cells. Transcription factor program enrichment analysis (via ChEA) of genes differentially upregulated in EBV^+^ B cells in the INCOV cohort.

Table S4: Pathway enrichment in EBV^+^ B cells. Pathway enrichment analysis (via GO:BP and KEGG) of genes differentially upregulated in EBV^+^ B cells in the INCOV cohort.

Table S5: Kinase enrichment in EBV^+^ B cells. Kinase enrichment analysis (via ARCHS4) of genes differentially upregulated in EBV^+^ B cells in the INCOV cohort.

Table S6: Differentially expressed genes in EBV^+^ B cells. Differential gene expression analysis between EBV^+^ and EBV^-^ B cells in the HAARVI cohort.

Table S7: Transcription factor enrichment in EBV^+^ B cells. Transcription factor program enrichment analysis (via ChEA) of genes differentially upregulated in EBV^+^ B cells in the HAARVI cohort.

Table S8: Pathway enrichment in EBV^+^ B cells. Pathway enrichment analysis (via GO:BP and KEGG) of genes differentially upregulated in EBV^+^ B cells in the HAARVI cohort.

Table S9: Kinase enrichment in EBV^+^ B cells. Kinase enrichment analysis (via ARCHS4) of genes differentially upregulated in EBV^+^ B cells in the HAARVI cohort.

Table S10: Enrichment of disease associated genes in EBV^+^ B cells. Enrichment analysis of disease associated genes (via GWAS Catalog and Orphanet) in EBV^+^ B cells in the INCOV cohort.

Table S11: Shared genes between EBV^+^ B cell states. Overlapping genes between upregulated genes in EBV^+^ naive B cells, EBV^+^ memory B cells, and EBV^+^ plasmablasts in the INCOV cohort.

Table S12: Differential metabolic flux reactions in EBV^+^ memory B cells. Differential activity analysis of metabolic flux reactions in EBV^+^ B cells in the INCOV cohort.

Table S13: Serum metabolite correlates of EBV^+^ B cell repertoires. Correlations (Pearson’s method) between metabolite plasma levels and the proportion of a patient’s B cell compartment that is EBV^+^ in the INCOV cohort.

Table S14: Cell-cell communication of EBV^+^ B cells with EBV^-^ B cells and CD4^+^ T cells. Cell-cell communication interaction scores (computed via CellPhoneDB^106^) between EBV^+^ B cells and EBV^-^ B cells and CD4^+^ T cells in the INCOV cohort.

Table S15: COVID-19 EBV reprogramming signatures in naïve and memory B cells. Gene sets defined in this work that are enriched in EBV reprogrammed naïve and memory B cells from patients with COVID-19 (INCOV cohort).

